# Concordant dynamic changes of global network properties in the frontoparietal and limbic compartments: an EEG study

**DOI:** 10.1101/2023.10.05.561067

**Authors:** Tien-Wen Lee, Gerald Tramontano, Clay Hinrichs

**Author notes:** Corresponding author at The NeuroCognitive Institute, New Jersey, US, Address: 111 Howard Blvd, Suite 204, Mt. Arlington, NJ 07856, Web: http://neuroci.com. Email addresses for the other author: G Tramontano, C Hinrichs.

## Abstract

**Introduction:** Despite its complexity, deciphering nodal interaction is imperative to understanding a neural network. Network interaction is an even more complicated topic that must be addressed. This study aimed to examine the relationship between the brain waves of two canonical brain structures, i.e., the frontoparietal and limbic compartments, during a resting state.

**Methods:** Electroencephalography (EEG) of 51 subjects in eye-closed condition was analyzed, and the eLORETA method was applied to convert the signals from the scalp to the brain. By way of community detection, representative neural nodes and the associated mean activities were retrieved. Total and lagged coherences were computed to indicate functional connectivity between those neural nodes. Two global network properties were elucidated based on the connectivity measures, i.e., global efficiency and mean functional connectivity strength. The temporal correlation of the global network indices between the two studied networks was explored.

**Results:** It was found that there was a significant trend of positive correlation across the four metrics (lagged vs. total coherence x global efficiency vs. average connectivity). In other words, when the neural interaction in the FP network was stronger, so did that in the limbic network, and vice versa. Notably, the above interaction was not spectrally specific and only existed at a finer temporal scale (under hundreds of milliseconds level).

**Conclusion:** The concordant change in network properties indicates an intricate balance between FP and LM compartments. Possible mechanisms and implications for the findings are discussed.

## Introduction

At a large-scale level, it is generally believed that brain functions are mediated by pertinent neural networks that may comprise several distinct and even distant brain regions (Pessoa, 2014). For instance, attention and working memory entail orchestrated computation in the frontoparietal (FP) matrix, and emotion, neuroendocrine, and vegetative regulation require coupled activations in the limbic (LM) system (Mega et al., 1997). Delineation of the network associated with specific cognition/function/behavior is one of the principal missions of brain research that can be accessed from various spectral, spatial, and temporal domains and scales. Networks are often studied in an isolated manner. Two classic examples are the FP and the default-mode networks (DMN) that have been corroborated by abundant literature (Fox et al., 2005; Kawasaki et al., 2010; Lee, 2016; Lee and Xue, 2018a; Raichle and Snyder, 2007; Vossel et al., 2014). Visual working memory requires coordination between frontal theta and parietal alpha activities (Kawasaki et al., 2010). DMN stands out in the “task-negative” condition (Fox et al., 2005; Raichle and Snyder, 2007). However, some essential neuropsychological functions involve the participation of more than one network (i.e., network interaction), such as emotion regulation which is substantiated by ameliorating LM activity through mobilizing the FP resource (Lee and Xue, 2018b), and directed attention which is characterized by enhancing perceptual processing via an influence of the attention network (Nebel et al., 2005).

Brain networks are complicated. The above examples are far from enough to address their innate complexity, not to mention their interactions. Exploring the interaction between brain networks has started to gain attention in the neuroimaging field (Hugdahl et al., 2019). Earlier research tended to probe this challenging issue from a hierarchical standpoint (Hilgetag et al., 2000; Sales-Pardo et al., 2007). Recently, Pascual-Marqui and Biscay-Lirio extended singular value decomposition to elucidate senders, hubs, and receivers from multiple time-lagged signals (Pascual-Marqui and Biscay-Lirio, 2010). The interaction between senders and receivers was mediated through the hubs, and they were all situated at the same level/hierarchy. By identifying microstates (networks) and treating each state as a whole, i.e., each microstate/network was as taken as a univariate, it was possible to study the directional influence of the microstates by way of, say, isolated coherence (Pascual-Marqui et al., 2014). The motivation behind the latter approach is of particular interest since it avoided studying the cumbersome node-to-node interactions between the studied networks and simplified the bi-network interaction issue to a bi-variate problem (no matter how many nodes reside in the network).

This research planned to investigate the interaction between two canonical networks, i.e., FP and LM, by electroencephalography (EEG). Although previous research has probed this issue using the emotion regulation paradigm (Lee and Xue, 2018b), their communication at a resting state still needs to be discovered. The central nervous system is constantly active, and the neural informatics embedded in the resting brain may contain a replica in the activated state, may affect the evoked responses, and are clustered into prominent modular structures (He, 2013; Lee et al., 2011c; Lee et al., 2014; Luczak et al., 2009; Moussa et al., 2012; Seeley et al., 2007; Tsodyks et al., 1999). There are opposing speculations about the network interaction between the two big compartments in the resting state. On the one hand, research from emotion–cognition and task-positive/-negative perspectives tend to infer that the FP and LM systems are antagonistic to some extent (Lee et al., 2013; Lee and Xue, 2018b). On the other hand, synergistic activations in the LP and LM networks have been noticed in specific scenarios, such as autobiographical planning and social working memory (Gerlach et al., 2011; Meyer et al., 2012; Spreng et al., 2010; Spreng, 2012). In addition, cortical oscillation in a resting state has long been suggested to have a thalamic origin, though not exclusively (Fuentealba and Steriade, 2005). If a third party modulated the two networks simultaneously, their interaction would be concordant, not discordant. The above ambiguity warrants empirical research to clarify.

To substantiate brain-based exploration, exact low-resolution brain electromagnetic tomography (eLORETA) was exploited to project EEG data from the scalp to the gray matter voxels of a template brain (Jurcak et al., 2007; Pascual-Marqui, 2007a). Data reduction based on community detection was applied to retrieve representative neural nodes and the associated brain dynamics in the frontal, parietal, and LM regions (note: we used compartment/region and network to denote the informatics before and after dimension reduction henceforth) (Lee and Xue, 2018a; Lee and Tramontano, 2021). These abridged brain signal traces formed the foundation for subsequent network interaction analyses, which comprised two steps. First, the signal traces were cut into shorter segments (0.4, 0.8, and 1.2 sec), and the network property of each EEG segment of FP and LM was constructed. Here, the “network property” is a univariate index that indicates a global index of a network, like a method initiated by Pascual-Marqui et al. introduced above (Pascual-Marqui et al., 2014). Second, the dynamic changes in network properties were correlated between the FP and LM networks. The null hypothesis is that the network properties of FP and LM networks fluctuate independently of each other.

## Materials and Methods

### EEG dataset and preprocessing

A public-released EEG dataset (BCI2000 instrumentation system) was used in this study (Goldberger et al., 2000; Schalk et al., 2004). The (resting) EEG traces contain 64 channels and last for 1 min, sampled at 160 Hz. Only the eyes-close resting condition (fewer electro-ocular artifacts) was included in the analysis to retain as much data as possible. The 64 channels covered the scalp in an approximately uniformly distributed manner following the 10-10 system. Software EEGLAB was applied to edit the EEG traces (Delorme and Makeig, 2004). The preprocessing steps included a band-pass filter (1–50 Hz), automatic artifact removal (AAR toolbox; ocular and electromyogenic artifacts), and manual elimination of the remaining noisy portions. The obtained clean EEGs were cut into 0.4 sec (64=2^6 data points), 0.8 sec (128=2^7 points), and 1.2 sec (192 points) segments with 50 percent sliding overlap, which were then exported to eLORETA. We investigated the time scales below 1.2 sec because of the consideration of correlation computation. Power analysis suggested that at alpha 0.05 and power 0.80, 80 points are the minimal required number for trustworthy correlation analysis (https://www.psychologie.hhu.de/arbeitsgruppen/allgemeine-psychologie-und-arbei tspsychologie/gpower). Compatible with the above estimate, a 1.2-sec segment with 50% overlap would yield 99 pieces at the maximum (60/1.2*2-1). After trimming EEG traces, the deletion of 15 percent of the data would still suffice for correlation analyses of the two studied networks. Notably, the segments that intervened with deletion were excluded because of the consideration of valid spectral transformation.

The scalp-recorded EEG signals were converted to brain-based neural activities by eLORETA (i.e., topography to tomography) (Jurcak et al., 2007; Pascual-Marqui, 2007a). In brief, the eLORETA linear imaging method provides a weighted minimum norm inverse solution. The weights are adaptive to the data (data-dependent) and may accommodate measurement and biological noises. Any arbitrary point-test sources can be correctly computed with exact, zero-error localization. Based on the principles of linearity and superposition, eLORETA is suitable for delineating distributed electric sources in the brain, albeit with low spatial resolution. Like its earlier version, sLORETA, the regional inference is based on standardized current density.

### Data reduction based on community detection

The outputs of the eLORETA inverse solution, time series of current source density, were registered to 6,239 gray matter voxels based on a template brain. As the name “low resolution” indicates, the tomographic images are “smooth, “ meaning that neighboring voxels’ derived neural activities could be highly similar. With this regard, it is tempting to adopt a data reduction strategy to reduce the data dimension to facilitate disclosing the underlying network structure and property. We resorted to a community detection algorithm to achieve this goal (Lee and Xue, 2018a; Lee and Tramontano, 2021). Brodmann areas (BAs) were referred to identify voxels in the frontal, parietal, and LM regions. In this study, the LM compartment contained both limbic and paralimbic regions. It was divided into two sub-components, i.e., orbitofrontal-centered (paleocortical) and hippocampal-centered (archicortical) belts, according to Mega et al. (Mega et al., 1997). The frontal (BA 6, 8–10, 44–46), parietal (BA 7, 39, 40), orbitofrontal-centered LM (BA 11, 32, 38), and hippocampal-centered LM regions (BA 23–25, 27–31, 33–36) contain 1429, 1107, 555, and 713 voxels, respectively.

Modules or communities are groups of nodes within a network that are more densely connected to one another than to other nodes. A correlation matrix was constructed for each of the four brain regions (with each element the correlation coefficient between voxel pair) based on the eLORETA outputs. Within this context, the neural nodes are the voxels, and correlation coefficients define the connectivity strengths between nodes. It is straightforward to show that the voxels with similar EEG dynamics would possess higher functional connectivity and tend to aggregate as a community. Dimensionality reduction through community detection based on correlation strengths has been successfully applied in high-dimensional neuroimaging data (Lee and Xue, 2018a; Lee and Tramontano, 2021). Although there are plenty of data reduction schemes, the above approach fits nicely with the rationale behind data reduction in neuroimaging research – the voxels with higher signal similarities are grouped. In contrast to Principal Component Analysis or Independent Component Analysis, the community-detection-based data reduction strategy avoids any spatial transformation. It maintains the measured “raw data” at the pertinent neural nodes.

The Louvain method was employed to partition each of the four regions into composite modules through optimization of Newman’s modularity metric Q (Blondel et al., 2008; Leicht and Newman, 2008; Rubinov and Sporns, 2011), in which 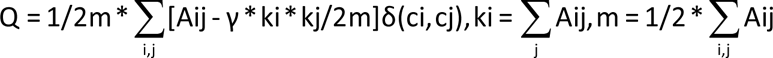, Aij is the connectivity strength between nodes i and j, ci is the community that node i belongs to, δ is the delta function (if ci=cj, delta=1, otherwise 0), and γ is the resolution factor (default=1). The above computations of community detection were performed using the Brain Connectivity Toolbox (http://www.brain-connectivity-toolbox.net) (Rubinov and Sporns, 2011). The mathematical platform MATLAB was used to execute the EEGLAB, Brain Connectivity Toolbox, and statistical analyses (Mathworks, Natick, MA, USA). We took an average time course for each module (neural node) as the representative brain dynamics, forming the basis of network property estimation through coherence and global efficiency computation (depicted below). The retrieved network nodes in the frontal and parietal regions were assembled to construct the FP network, and so were the nodes in the two LM sub-regions to form the LM network.

### Network property: global efficiency and mean coherence for each EEG segment

We resorted to coherence as the functional connectivity for network property analysis. The spectra of delta (1.5–4 Hz), theta (4–8 Hz), alpha (8–12 Hz), beta1 (12–18 Hz), beta2 (18–22 Hz), beta3 (22–30 Hz), and broadband (1.5–30 Hz) were explored (Lee and Tramontano). Coherence is commonly used to estimate functional connectivity between two signal series in the frequency domain (Pfurtscheller and Andrew, 1999), defined as Cxy(f) = |Gxy(f)|^2/(Gxx(f)*Gyy(f)), where f is the studied frequency band, Gxy(f) the cross-spectral density between two signal series x(t) and y(t), and Gxx(f) and Gyy(f) the auto-spectral density of x(t) and y(t), respectively. Two coherence indices were investigated in this research, i.e., total coherence and lagged coherence. Total coherence is the sum of the lagged and instantaneous dependence. The latter could contain confounds from the “non-physiological contribution due to volume conduction and low spatial resolution” (Pascual-Marqui, 2007b). After the Fourier Transform of the EEG signal (there shall be real and imagery components), the instantaneous coherence considers the real part of the Hermitian covariance matrix at the defined frequency. The exact formula of the two coherence indices can be referred to (Pascual-Marqui, 2007b). We expected the results derived from lagged coherence to be more conservative and reliable than total coherence. Given that total coherence is still used in EEG research, we present both for comparison but focus on the results derived from lagged coherence.

Since the main interest of this research is about the characteristics of a network, not a particular neural node, we resorted to the measure of network integration (in opposition to the segregation index of a specific neural node) using a graph theoretical approach, namely global efficiency. In detail, we took the inverse of coherence as the “distance” between two network nodes (the higher the coherence, the shorter the distance). The shortest weighted path length between any two nodes was thus delineated, and hence the weighted global efficiency (i.e., the average of the inverse of the shortest path lengths between all the pairs of network nodes); the formula can be referred to Table A1 in (Rubinov and Sporns, 2010). In addition, the mean coherence of all the possible connections (excluding self-connection, which is always 1) was also calculated to represent the degree of global interaction in the FP and LM networks. In sum, four network properties are derived from bi-variate coherence metrics, i.e., lagged vs. total coherence x global efficiency vs. average coherence. The streamlining of the analytic flow is illustrated in Figure 1.

**Fig. 1.**
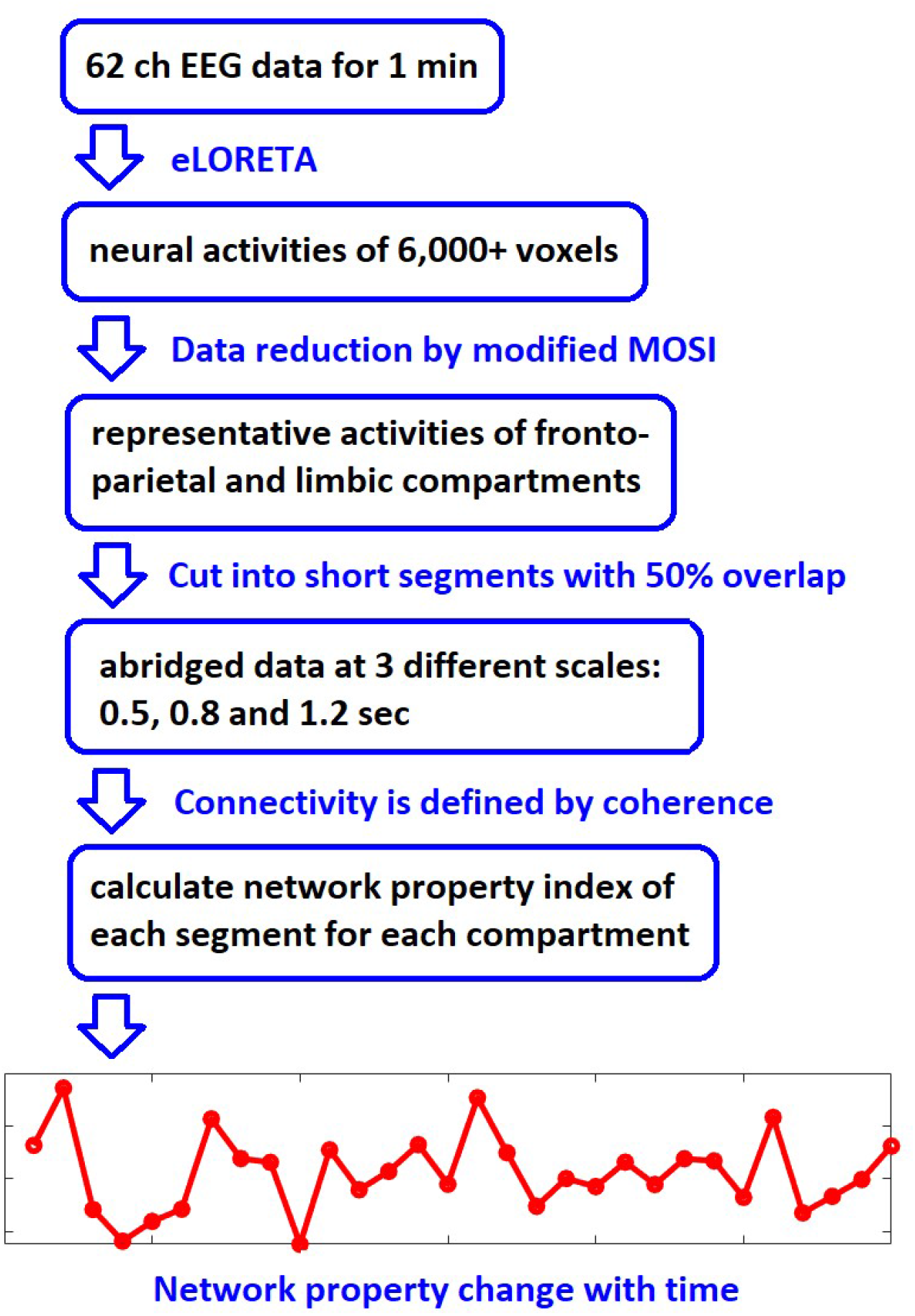
Flowchart of the analytic flow to derive the global network properties.

### Examination of the concordance of network property between the FP and LM networks

Four network property indices were computed for each EEG segment for the FP and LM networks, respectively, as described above. Pearson correlation analyses were performed for each subject to investigate the concordance of network property change with time. We used geometric mean to derive the group-level *p-value*: Suppose there are k subjects, with the *p-values* of correlation p1, p2, …, and pk. The group level *p-value* is calculated as (p1*p2*…pk)^(1/k). The null hypothesis was examined by this group-level *p-value* <= 0.01.

## Results

We selected 51 out of 109 subjects and imported EEG data from 62 electrodes into the analyses (T9 and T10 were excluded). Quality check of the dataset is described in Part I of the **Supplementary Material**. After careful trimming procedures, the average length of EEG traces is 59.1 sec (SD 0.78; min 54.48 sec and max 60 sec). In the main text, we only report results from EEG signals segmented at the scales of 0.4 sec and 0.8 sec. The 1.2-sec counterparts are generally insignificant; see Part II in the **Supplementary Material**.

### Data reduction through modular analysis

The Louvain algorithm partitioned the voxels in the frontal, parietal, orbitofrontal-centered, and hippocampal-centered LM regions into 4.3 (SD 0.53, 3–5), 3.2 (SD 0.42, 3–4), 3.6 (SD 0.49, 3–4) and 3.1 (SD 0.35, 3–4) communities. The representative temporal dynamics of each cluster were retrieved by averaging the temporal dynamics of the voxels belonging to the same cluster.

### Network property I: global efficiency based on lagged and total coherences

The global efficiency in the FP and LM networks are summarized in the upper part of Table 1 (EEG traces segmented at 0.4 sec) and Table 2 (EEG traces segmented at 0.8 sec). Unsurprisingly, the mean values of global efficiency based on the total coherence are higher than those of the lagged coherence. As to network interaction, at a finer time scale (0.4 sec), there is a significant and spectrum-unspecific trend that the temporal changes of the global efficiency correlated positively between FP and LM networks, see Table 1 (lower part). The trend is less robust at coarser temporal resolution (0.8 sec); see Table 2 (lower part).

**Table 1.** Global efficiency (upper) and its correlation (lower) between the frontoparietal and limbic networks (based on total and lagged coherence). EEG trace is segmented at the scale of 0.4 sec.

| **Spectra** | delta | theta | alpha | beta1 | beta2 | beta3 | 1.5–30 Hz |
| --- | --- | --- | --- | --- | --- | --- | --- |
| **Global Efficiency: Mean (SD)** |  |  |  |  |  |  |  |
| **Based on total coh** |  |  |  |  |  |  |  |
| Frontoparietal | 0.505 (0.08) | 0.470 (0.07) | 0.393 (0.05) | 0.385 (0.06) | 0.384 (0.07) | 0.334 (0.05) | 0.192 (0.04) |
| Limbic | 0.488 (0.07) | 0.459 (0.06) | 0.389 (0.06) | 0.383 (0.06) | 0.378 (0.06) | 0.332 (0.05) | 0.211 (0.04) |
| **Based on lagged coh** |  |  |  |  |  |  |  |
| Frontoparietal | 0.311 (0.05) | 0.296 (0.04) | 0.238 (0.04) | 0.235 (0.04) | 0.235 (0.04) | 0.197 (0.03) | 0.087 (0.02) |
| Limbic | 0.281 (0.04) | 0.270 (0.04) | 0.213 (0.03) | 0.210 (0.03) | 0.211 (0.03) | 0.174 (0.03) | 0.075 (0.01) |
| **Network Interaction: Correlation** |  |  |  |  |  |  |  |
| **Based on total coh** |  |  |  |  |  |  |  |
| Correlation Coeff | 0.284 (0.14) | 0.267 (0.13) | 0.287 (0.14) | 0.293 (0.15) | 0.326 (0.13) | 0.313 (0.13) | 0.375 (0.12) |
| Group *p*-val | 1.256E-08 | 1.293E-07 | 7.001E-09 | 2.397E-09 | 1.845E-10 | 8.522E-010 | 7.926E-13 |
| **Based on lagged coh** |  |  |  |  |  |  |  |
| Correlation Coeff | 0.254 (0.11) | 0.199 (0.11) | 0.200 (0.10) | 0.214 (0.09) | 0.227 (0.09) | 0.214 (0.11) | 0.230 (0.10) |
| Group *p*-val | 3.704E-07 | 4.750E-05 | 2.447E-05 | 1.536E-05 | 2.954E-06 | 2.383E-05 | 8.003E-06 |

Note: The group *p*-value is derived by taking the geometric mean of individual *p*-values. Abbreviations: Coh, coherence; Coeff, coefficient.

**Table 2.** Global efficiency (upper) and its correlation (lower) between the frontoparietal and limbic networks (based on total and lagged coherence). EEG trace is segmented at the scale of 0.8 sec.

| **Spectra** | delta | theta | alpha | beta1 | beta2 | beta3 | 1.5–30 Hz |
| --- | --- | --- | --- | --- | --- | --- | --- |
| **Global Efficiency: Mean (SD)** |  |  |  |  |  |  |  |
| **Based on total coh** |  |  |  |  |  |  |  |
| Frontoparietal | 0.415 (0.06) | 0.335 (0.04) | 0.300 (0.04) | 0.294 (0.05) | 0.293 (0.05) | 0.244 (0.04) | 0.135 (0.03) |
| Limbic | 0.395 (0.06) | 0.331 (0.05) | 0.305 (0.05) | 0.302 (0.05) | 0.296 (0.05) | 0.254 (0.04) | 0.164 (0.04) |
| **Based on lagged coh** |  |  |  |  |  |  |  |
| Frontoparietal | 0.245 (0.04) | 0.196 (0.03) | 0.169 (0.02) | 0.167 (0.03) | 0.166 (0.03) | 0.131 (0.02) | 0.051 (0.01) |
| Limbic | 0.209 (0.04) | 0.170 (0.03) | 0.149 (0.02) | 0.149 (0.02) | 0.147 (0.02) | 0.114 (0.02) | 0.042 (0.01) |
| **Network Interaction: Correlation** |  |  |  |  |  |  |  |
| **Based on total coh** |  |  |  |  |  |  |  |
| Correlation Coeff | 0.297 (0.15) | 0.256 (0.15) | 0.305 (0.14) | 0.289 (0.15) | 0.315 (0.14) | 0.294 (0.14) | 0.373 (0.15) |
| Group *p*-val | 2.988E-05 | 2.174E-04 | 1.891E-05 | 3.753E-05 | 1.426E-05 | 4.550E-05 | 2.337E-07 |
| **Based on lagged coh** |  |  |  |  |  |  |  |
| Correlation Coeff | 0.226 (0.14) | 0.173 (0.13) | 0.199 (0.14) | 0.179 (0.13) | 0.202 (0.14) | 0.173 (0.13) | 0.227 (0.12) |
| Group *p*-val | 1.033E-03 | 1.033E-02 | 2.962E-03 | 6.926E-03 | 2.730E-03 | 9.459E-03 | 1.758E-03 |

Note: The group *p*-value is derived by taking the geometric mean of individual *p*-values. Abbreviations: Coh, coherence; Coeff, coefficient.

### Network property II: mean connectivity strength based on lagged and total coherences

The mean connectivity strengths (i.e., mean coherence) in the FP and LM networks are summarized in the upper part of Table 3 (EEG traces segmented at 0.4 sec) and Table 4 (EEG traces segmented at 0.8 sec). The same as in the previous section, it is noticed that the mean values of the total coherence are always higher than those of the lagged coherence. As to network interaction, again, there is a significant and spectrum-unspecific trend that the temporal changes in the mean connectivity correlated positively between FP and LM networks, summarized in Tables 3 and 4 (lower part). The trend is significant on a finer scale and less prominent on a coarser temporal scale.

**Table 3.** Average connectivity (upper) and its correlation between (lower) the frontoparietal and limbic networks across 7 spectra (based on total and lagged coherence). EEG trace is segmented at the scale of 0.4 sec.

| Spectra | delta | theta | alpha | beta1 | beta2 | beta3 | 1.5–30 Hz |
| --- | --- | --- | --- | --- | --- | --- | --- |
| **Average Connectivity: Mean (SD)** |  |  |  |  |  |  |  |
| **Based on total coh** |  |  |  |  |  |  |  |
| Frontoparietal | 0.493 (0.08) | 0.453 (0.07) | 0.368 (0.05) | 0.359 (0.06) | 0.359 (0.07) | 0.305 (0.05) | 0.167 (0.04) |
| Limbic | 0.477 (0.07) | 0.443 (0.06) | 0.367 (0.05) | 0.360 (0.06) | 0.356 (0.06) | 0.307 (0.05) | 0.188 (0.04) |
| **Based on lagged coh** |  |  |  |  |  |  |  |
| Frontoparietal | 0.248 (0.04) | 0.232 (0.04) | 0.182 (0.03) | 0.179 (0.03) | 0.180 (0.03) | 0.148 (0.02) | 0.064 (0.01) |
| Limbic | 0.224 (0.04) | 0.213 (0.03) | 0.164 (0.02) | 0.162 (0.03) | 0.163 (0.03) | 0.133 (0.02) | 0.060 (0.01) |
| **Network Interaction: Correlation** |  |  |  |  |  |  |  |
| **Based on lagged coh** |  |  |  |  |  |  |  |
| Correlation Coeff | 0.287 (0.14) | 0.270 (0.12) | 0.288 (0.14) | 0.291 (0.15) | 0.321 (0.13) | 0.307 (0.12) | 0.375 (0.12) |
| Group *p*-val | 1.285E-08 | 1.492E-07 | 1.096E-08 | 5.367E-09 | 5.194E-10 | 2.821E-09 | 8.864E-13 |
| **Based on total coh** |  |  |  |  |  |  |  |
| Correlation Coeff | 0.245 (0.13) | 0.192 (0.11) | 0.200 (0.13) | 0.213 (0.12) | 0.223 (0.13) | 0.214 (0.11) | 0.237 (0.11) |
| Group *p*-val | 1.015E-06 | 9.500E-05 | 3.098E-05 | 1.947E-05 | 3.952E-06 | 2.864E-05 | 4.307E-06 |

Note: The group *p*-value is derived by taking the geometric mean of individual *p*-values. Abbreviations: Coh, coherence; Coeff, coefficient.

**Table 4.** Average connectivity (upper) and its correlation between (lower) the frontoparietal and limbic networks across 7 spectra (based on total and lagged coherence). EEG trace is segmented at the scale of 0.8 sec.

| Spectra | delta | theta | alpha | beta1 | beta2 | beta3 | 1.5–30 Hz |
| --- | --- | --- | --- | --- | --- | --- | --- |
| **Average Connectivity: Mean (SD)** |  |  |  |  |  |  |  |
| **Based on total coh** |  |  |  |  |  |  |  |
| Frontoparietal | 0.392 (0.06) | 0.305 (0.04) | 0.270 (0.04) | 0.263 (0.04) | 0.263 (0.05) | 0.214 (0.04) | 0.115 (0.04) |
| Limbic | 0.375 (0.06) | 0.305 (0.05) | 0.279 (0.04) | 0.275 (0.05) | 0.270 (0.05) | 0.228 (0.04) | 0.143 (0.03) |
| **Based on lagged coh** |  |  |  |  |  |  |  |
| Frontoparietal | 0.189 (0.03) | 0.147 (0.02) | 0.126 (0.02) | 0.124 (0.02) | 0.124 (0.02) | 0.097 (0.01) | 0.037 (0.01) |
| Limbic | 0.162 (0.03) | 0.129 (0.02) | 0.113 (0.02) | 0.112 (0.02) | 0.111 (0.02) | 0.086 (0.01) | 0.032 (0.01) |
| **Network Interaction: Correlation** |  |  |  |  |  |  |  |
| **Based on lagged coh** |  |  |  |  |  |  |  |
| Correlation Coeff | 0.302 (0.14) | 0.255 (0.14) | 0.306 (0.14) | 0.283 (0.15) | 0.310 (0.14) | 0.292 (0.14) | 0.372 (0.15) |
| Group *p*-val | 3.025E-05 | 3.003E-04 | 1.987E-05 | 5.975E-05 | 2.162E-05 | 5.095E-05 | 2.402E-07 |
| **Based on total coh** |  |  |  |  |  |  |  |
| Correlation Coeff | 0.225 (0.14) | 0.177 (0.12) | 0.203 (0.14) | 0.180 (0.13) | 0.203 (0.14) | 0.175 (0.13) | 0.233 (0.12) |
| Group *p*-val | 1.242E-03 | 9.723E-03 | 2.473E-03 | 6.740E-03 | 2.644E-03 | 8.739E-03 | 1.514E-03 |

Note: The group *p*-value is derived by taking the geometric mean of individual *p*-values. Abbreviations: Coh, coherence; Coeff, coefficient.

The results derived from EEG traces segmented at 0.4, 0.8, and 1.2 sec altogether suggest that the interaction between FP and LM networks is temporal scale sensitive. It vacillated relatively fast, and the relationship was masked when the temporal window was lengthened to 0.8 sec or longer. The distributions of correlation coefficients of global efficiency and average coherence between the FP and LM networks across the three temporal scales are illustrated in Figure 2.

**Fig. 2.**
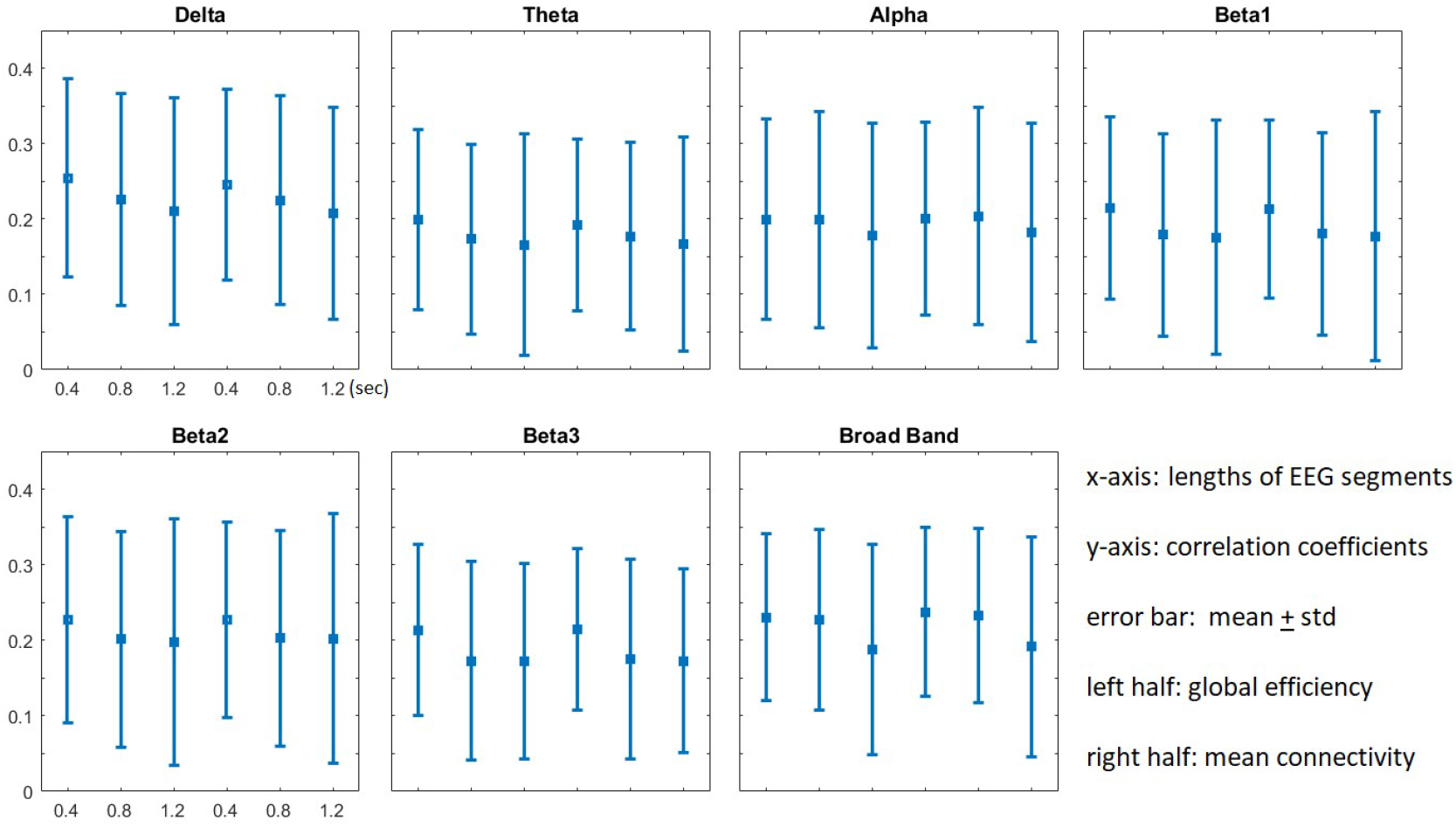
Plots of the correlation coefficients (Y-axis) of network properties between the two studied compartments across three conditions: EEG traces segmented at 0.4, 0.8, and 1.2 sec (labeled at the X-axis). The square mark and error bar indicates the mean value and one standard deviation (std), respectively. Left half for each subplot: correlation coefficients of global efficiency. Right half for each subplot: correlation coefficients of mean coherence. Broad Band indicates the spectrum 1.5–30 Hz.

## Discussion

In contrast to the neocortex, the LM system is phylogenetically more ancient. During an active engagement in psychological tasks, it has been demonstrated that the relationship between the FP and LM networks could be either antagonistic (e.g., in emotion regulation or a wide range of externally directed tasks) or synergistic (e.g., in autobiographical planning, simulated problem-solving, or social working memory) (Gerlach et al., 2011; Meyer et al., 2012; Spreng et al., 2010; Spreng, 2012). Although evidence supports intense crosstalk between the FP and LM compartments under various contexts, their interaction in the resting state is largely unknown. We examined 51 EEG datasets recorded in an eye-closed condition. The scalp signals were projected to a template brain based on eLORETA. Data reduction procedures were adopted to convert a brain region with hundreds of voxels to several neural nodes (generally less than 6). Coherence (total and lagged) indicated functional connectivity between two neural nodes. Two network properties were explored, i.e., global efficiency and mean connectivity strength. Our results showed that during the resting state, the fluctuation of the two explored network properties was positively correlated between LP and LM. In other words, when the neural interaction in the FP network was more robust, so did that in the LM network, and vice versa. This pattern was temporal scale sensitive and consistently demonstrated across the spectra from delta to beta3. The lack of spectral specificity was verified in the broadband spectrum analysis. The null hypothesis is declined.

It is acknowledged that brain networks and network interaction are complicated. To name a few: Distinct patterns of theta/high gamma coupling were noticed across the human neocortex for different behavioral tasks (Canolty et al., 2006). Delta and alpha seemed to play different roles in the working memory exercise (Kawasaki et al., 2010). Incoherent brain signals could interact through amplitude envelope correlation (Bruns et al., 2000). Jirsa et al. summarized six possible forms of cross-frequency relationships in a neural network (Jirsa and Muller, 2013). Unlike previous studies that explored the information exchange between neural nodes within a pre-defined network (Kawasaki et al., 2010) or the spectral/hierarchical relationship within a brain region (Jirsa and Muller, 2013), this study focused on the interaction between outer cortex (FP mantle, excluding primary sensorimotor components) and inner cortex (LM system, including paralimbic regions). These two compartments are relatively separable, and the neural substrates can be delineated by cytoarchitecture or neuroimaging methods (Lee et al., 2014; Mega et al., 1997). Perhaps because of the relative independence, their interaction was reported to be context dependent (synergistic or antagonistic), unlike the behaviors of neural nodes in a network that are hardwired by white matter bundles. For example, frontal and parietal activations/deactivations are generally consistent (Lee and Xue, 2018b; Wager and Smith, 2003). Then, what are the potential pathways that mediate this compartmental-level interaction?

Firstly, it was shown that DMN might carry the information flow of the LM system (Lee and Xue, 2018a). The several transmodal regions of DMN in the outer cortex are plausible candidates, such as the middle temporal, inferior parietal, and (anti-correlated) prefrontal cortices (Mesulam, 1998). Secondly, the neural substrates for emotion regulation could also provide venues to substantiate the interaction (Kringelbach and Rolls, 2004; Lee and Xue, 2018b). However, the observed concordance between the FP and LM networks appeared to be present across all the spectra. This spectral non-specificity was rare in the neuroscientific literature regarding cortico-cortical communication (Klimesch, 1999; Sauseng et al., 2002). An earlier work of imaging genetics by the author Lee might hint at this discovery. It was found that different genotypes of N-methyl D-aspartate receptor (NMDAR; a ligand-gated ionotropic glutamate receptor) could impose differential global influence on the EEG over broad spectra (Lee et al., 2011a). NMDAR plays essential roles in synaptic plasticity, learning, and memory (Pagano et al., 2021) and is necessary for the normal functioning of thalamo-cortico-thalamic and thalamo-cortico-striatal circuits (Amat-Foraster et al., 2019; Carli and Invernizzi, 2014; McCormick, 2002). Thalamic projection to the cortex is widespread, covering both FP and LM compartments, and may affect cortical oscillation across different spectra (e.g., from delta to beta bands) (Basha et al., 2014; Fuentealba and Steriade, 2005; Halgren et al., 2019; Steriade et al., 1993). Given that there lacks an informational “highway” between the FP and LM compartments (e.g., like the longitudinal fasciculi and cingulum), the thalamus is an ideal candidate to moderate the compartment-level interaction given its broad projection, reliance on intricate NMDAR system for ordinary physiology, and its well-acknowledged roles in the cortical oscillations (informational pathway contrasts with modulating analog, nicely summarized in (Garcia-Toro et al., 2001)). Other excitatory and inhibitory neuromodulating pathways are also worthy of consideration, such as serotonergic and dopaminergic systems (Lee et al., 2011b; Lee et al., 2012), which also affect a large portion of the brain and have been implicated in several neuropsychiatric conditions that are characterized by FP and LM dysregulations (Carli and Invernizzi, 2014). The above accounts (cortical and subcortical) are not exclusive to each other and may operate together to fulfill the FP and LM communication.

May our results illuminate the underlying mechanisms of FP and LM interaction? Let us assume a simplified scenario here: excitatory–excitatory, excitatory–inhibitory, inhibitory–excitatory, and inhibitory–inhibitory. It is noteworthy that our findings negate the excitatory–inhibitory and inhibitory–excitatory ones but do not exclude the possibility of excitatory–excitatory and inhibitory–inhibitory counterparts. Evidence supports that the neural patterns in the active state may re-instantiate themselves in the resting state (Luczak et al., 2009; Tsodyks et al., 1999). The resting brain is not idling, in which neural plasticity and learning processes may occur. The inferred excitatory–excitatory and inhibitory–inhibitory patterns may accommodate synergistic and antagonistic circumstances of FP and LM interaction, respectively. The excitatory–excitatory indicates a “concordant” positive boost to each other and might happen in the milieu of FP and LM collaboration. The inhibitory–inhibitory implies a “concordant” offset to each other in the FP and LM antagonism. Overall, our findings suggest an intricate balance between FP and LM compartments (Lee, 2016; Lee and Xue, 2018c). This dynamic balance is so fundamental that it is maintained across different spectra and achieved promptly – at hundreds of milliseconds. Since the sampling rate of the EEG dataset was 160 points per sec, it would be interesting to probe the temporal scale issue by analyzing the EEG dataset collected at a higher speed to detect the temporal limit of the dynamic equilibrium. The findings also imply that other neuroimaging tools at a slower acquisition rate, such as functional magnetic resonance imaging, are likely to yield false negative results.

It is noteworthy that false positive bias is a serious issue when using coherence to indicate inter-nodal connectivity in EEG research. Conventional (total) coherence is prone to be confounded by instantaneous, non-physiological noises that exert influence over several channels simultaneously due to volume conduction and low spatial resolution (Pascual-Marqui, 2007b). To tackle this problem of exaggerated connectivity, lagged coherence was chosen as the main index, which assures that confounding factors were removed considerably. The rationale is straightforward: unlike non-physiological noises, proper neural connectivity via axonal and synaptic conduction is inevitably accompanied by travel time and hence phase “lag” in the frequency domain. After Fourier transformation, the proper connectivity is reflected in the imagery part, in contrast to the noises mainly reflected in the real part (instantaneous coherence). Per our expectation, the results derived from lagged coherence are more conservative than total coherence, see Tables 1 to 4. Our conclusions were drawn from the results of lagged coherence, not total coherence. Average reference innate in LORETA computation may also help remove noises from distant sources if they result in equal amounts of artifacts to all the EEG channels.

Particular concern exists about the methodology used to investigate the interaction between large-scale networks. In this study, the FP and LM cortices are arguably the two largest structures in the human cerebral cortex. Hence, the possible number of node-to-node interactions is enormous. To disclose their interaction, it is desirable to reduce the data dimension and treat the network as a whole (Pascual-Marqui et al., 2014). We achieve this goal in two steps. First, through a community detection algorithm using a correlation structure, the high dimensional data was abridged to several neural nodes and representative temporal dynamics. This method is of interest because it only requires one critical criterion (constraint): the regional voxels sharing similar information (indicated by higher correlation coefficients) are grouped (Lee and Xue, 2018a). Based on the same rationale, Lee and Tramontano recently invented “MOSI” which enables multi-resolution data reduction tailored for functional magnetic resonance imaging research (Lee and Tramontano, 2021). The output of eLORETA is inherently smooth, partly because of the limited electrode number and partly because of the similarity in signals of adjacent electrodes—precisely the situation in which compression of highly co-linear signals is justified. To the authors’ best knowledge, this is the first time eLORETA and community detection were combined to explore network issues. Secondly, we resorted to a graph theoretical approach to investigate (within-)network integration by global efficiency and the average strength of functional connectivity. This simplified scheme treated the characteristic behavior of a network as univariate and meanwhile eschewed handling the massive number of voxel pairs in the FP and LM compartments. Furthermore, it allows tracing the network property change with time. It is possible to apply multivariate methods to investigate the dependence or cross-spectral relationship between two (or more) networks (Pascual-Marqui et al., 2016; Pascual-Marqui, 2007a). However, these methods assume the EEG signal’s stationarity and require tens of EEG segments to construct the covariance matrix in the spectral domains, thus prohibiting it from investigating temporal vacillation of network property, as demonstrated in this study. Nevertheless, these inspiring methods in EEG research have yet to obtain enough attention and may provide novel insight into network interaction issues.

## Conclusion

The interaction between neural networks is an essential neuroscience topic that needs to be addressed more. This EEG study explored the relationship between two canonical brain structures, i.e., the frontoparietal and limbic compartments, during the resting state. The temporal correlation of the global network property changes unveiled a close coupling. The findings implicate an intricate and rapid balance between FP and LM compartments, which are present in all the studied spectra. In addition to several cortical hubs known to regulate the limbic system, the between-network concordance could also be mediated by the subcortical structures that project widely to the cortex, such as the thalamus, reticular formation, or neuromodulatory pathways. The synchronization in network properties could be one of the mechanisms underlying the communication between the frontoparietal and limbic compartments.

## Authors Contributions

TW Lee, G Tramontano, and C Hinrichs contributed intellectually to this work. TW Lee carried out the analysis and wrote the first draft. All authors reviewed the manuscript.

## Acknowledgments

This work was supported by NeuroCognitive Institute (NCI) and NCI Clinical Research Foundation Inc.

## Statements and Declarations

TW Lee, G Tramontano, and C Hinrichs declare no conflicts of interest.

## Compliance with ethical standards

The authors of this article carried out no animal or human studies. This research analyzed a publicly released dataset. The authors assert that all procedures contributing to this work comply with the ethical standards of the relevant national and institutional committees on human experimentation and with the Helsinki Declaration of 1975, as revised in 2008.

## Supplementary Materials

The **supplementary materials** comprise two parts. The first part is the quality assessment of data quality. The second part explores the issues concerning the temporal scale (1.2 sec) of the network interaction between frontoparietal (FP) and limbic (LM) compartments.

### I. Quality Assessment

The first part of the **supplementary materials** is about the data quality assessment. This study used a publicly released EEG dataset that was created and contributed to PhysioNet by the developers of the BCI2000 instrumentation system (Schalk et al., 2004). The dataset was collected from 109 volunteers and was intended for “EEG Motor Movement/Imagery” purposes. There were 14 experimental runs for each subject (https://archive.physionet.org/pn4/eegmmidb/). The second run (Baseline, eyes closed) was used in this research. We selected 51 out of the 109 subjects, highlighted in yellow in Column 2 of Table 1. The third column lists the reasons why some subjects were excluded from the analysis.

We noticed a faulty channel T10 which carried a systemic error to nearly every single subject (this was also noticed by other researchers, say Pascual-Marqui et al., 2014), see Figure 1. We thus excluded T9 and T10 from the 64-channel montage. It is understandable that some “noise” may have limited spectral distribution and won’t impose too much trouble when it comes to spectral analysis. However, since “broadband power” is one of the main foci of this study, we excluded those subjects that showed inadequate quality or undulation of multiple channels unless the artifacts can be corrected, say AAR and ASR of EEGLAB.

**Table 1.** quality check of the dataset contributed by Schalk et al., 2004.

|  |  |  |
| --- | --- | --- |
| 'S001' | T10 |  |
| 'S002' | T10 | T7 bad -> interpolation |
| 'S003' | T10 | C4, Cp6 |
| 'S004' | T10 | undulation of several channels |
| 'S005' | T10 | Poor quality |
| 'S006' | T10 | Poor quality |
| 'S007' | T10 | Poor quality |
| 'S008' | T10 | T9 a little bit drop at 10 Hz |
| 'S009' | T10 | Poor quality |
| 'S010' | T10 | undulation of several channels |
| 'S011' | T10 | AF8, CP2 |
| 'S012' | T10 | Fp1, T9, T8, Tp8 |
| 'S013' | T10 | alpha slowing to theta range |
| 'S014' | T10 | Ft8, Tp8, T8 |
| 'S015' | T10 | Ft8, T8 |
| 'S016' | T10 | Ft8, F8 |
| 'S017' | T10 | Ft7, Ft8 |
| 'S018' | T10 | T9 |
| 'S019' | T10 | drowsy? |
| 'S020' | T10 |  |
| 'S021' | T10 | F1, C5 |
| 'S022' | T10 | 5 bad channels (Fp1, Fp2, Fpz are the same) |
| 'S023' | T10 | drowsy? |
| 'S024' | T10 | multiple bad channels by ASR and topo |
| 'S025' | T10 | C6, TP7 |
| 'S026' | T10 | Af8, Af7 |
| 'S027' | Tp8 | F6, Tp8 |
| 'S028' | T10 | F8 |
| 'S029' | T10 | T8 |
| 'S030' | T10 |  |
| 'S031' | T10 | T7, FT7 |
| 'S032' | T10 | P1 bad, noisy |
| 'S033' | T10 | F7, CP5 |
| 'S034' | T10 | FT7, F7, T9, TP7 |
| 'S035' | T10 |  |
| 'S036' | T10 |  |
| 'S037' | T10 | multiple bad channels by ASR and topo |
| 'S038' | T10 |  |
| 'S039' | T10 |  |
| 'S040' | T10 | undulation of several channels |
| 'S041' | T10 | F7, T8, T9 |
| 'S042' | T10 | undulation of several channels |
| 'S043' | T9 | undulation of several channels |
| 'S044' | T10 | T9 |
| 'S045' | T10 |  |
| 'S046' | T10 | undulation of several channels |
| 'S047' | T10 | EKGs that can be removed by ave ref |
| 'S048' | T10 | F5, F8 |
| 'S049' | T10 |  |
| 'S050' | T10 |  |
| 'S051' | T10 | FC1, C5, FT7, T7; |
| 'S052' | T10 | FC3, F3; undulation of several channels |
| 'S053' | T10 | P7 -> interpolation; a lot of EKG can be removed by AAR |
| 'S054' | T10 | OK |
| 'S055' | T10 |  |
| 'S056' | T10 | OK |
| 'S057' | T10 | C6 |
| 'S058' | T10 |  |
| 'S059' | T10 | AF4, T8 |
| 'S060' | T10 | T8, T9 |
| 'S061' | T10 |  |
| 'S062' | T10 | undulation of several channels |
| 'S063' | T10 | PO8, P6, P8 |
| 'S064' | T10 | FT8, T8, TP8; undulation of multiple channels |
| 'S065' | T10 | T8, FT8 |
| 'S066' | T10 | drowsy? |
| 'S067' | T10 | F6, T8, AF8; undulation of several channels |
| 'S068' | T10 | CP1 -> interpolation |
| 'S069' | T10 | giant undulation of all channels (from ref) |
| 'S070' | T10 | EKG, FT7, T7 |
| 'S071' | T10 | EKG |
| 'S072' | T10 | undulation of several channels |
| 'S073' | T10 | EMG: T8, T9 |
| 'S074' | T10 | Cz, CPz, C2 |
| 'S075' | T10 | FC4 |
| 'S076' | T10 | weird spectrum |
| 'S077' | T10 | F8, FC5, FC6, C6 |
| 'S078' | T10 | pulsation: Af7 |
| 'S079' | T10 | FP1, FC6 |
| 'S080' | T10 | F7, F8, FT7 |
| 'S081' | T10 | F8 |
| 'S082' | T10 | T9, P6 |
| 'S083' | T10 | C3, AF7 |
| 'S084' | T10 | undulation of several channels |
| 'S085' | T10 | AF8, F7 -> interpolation |
| 'S086' | T10 | multiple bad channels by ASR and topo |
| 'S087' | T10 | pulsation: Fp2 |
| 'S088' | T10 | AF3, AF4, AF8 |
| 'S089' | T10 | F8, T7, F7 |
| 'S090' | T10 | T7, TP7 |
| 'S091' | T10 |  |
| 'S092' | T10 | undulation of several channels |
| 'S093' | T10 | undulation of several channels |
| 'S094' | T10 | EMG: T7 |
| 'S095' | T10 | C2 -> interpolation |
| 'S096' | T10 | Poor quality |
| 'S097' | T10 | TP7, AF7 |
| 'S098' | T10 | EMG; undulation of several channels |
| 'S099' | T10 |  |
| 'S100' | T10 | Fc5, T8, CP2 |
| 'S101' | T10 | multiple bad channels by ASR and topo |
| 'S102' | T10 | multiple bad channels by ASR and topo |
| 'S103' | T10 | large segments of artifact data |
| 'S104' | T10 | there are weird segments |
| 'S105' | T10 | FC1, FC3, FC5 |
| 'S106' | T10 | T9, EMG Af7, Fp1 |
| 'S107' | T10 | T8, T9 |
| 'S108' | T10 |  |
| 'S109' | T10 | C6, F4 |

**Figure 1.**
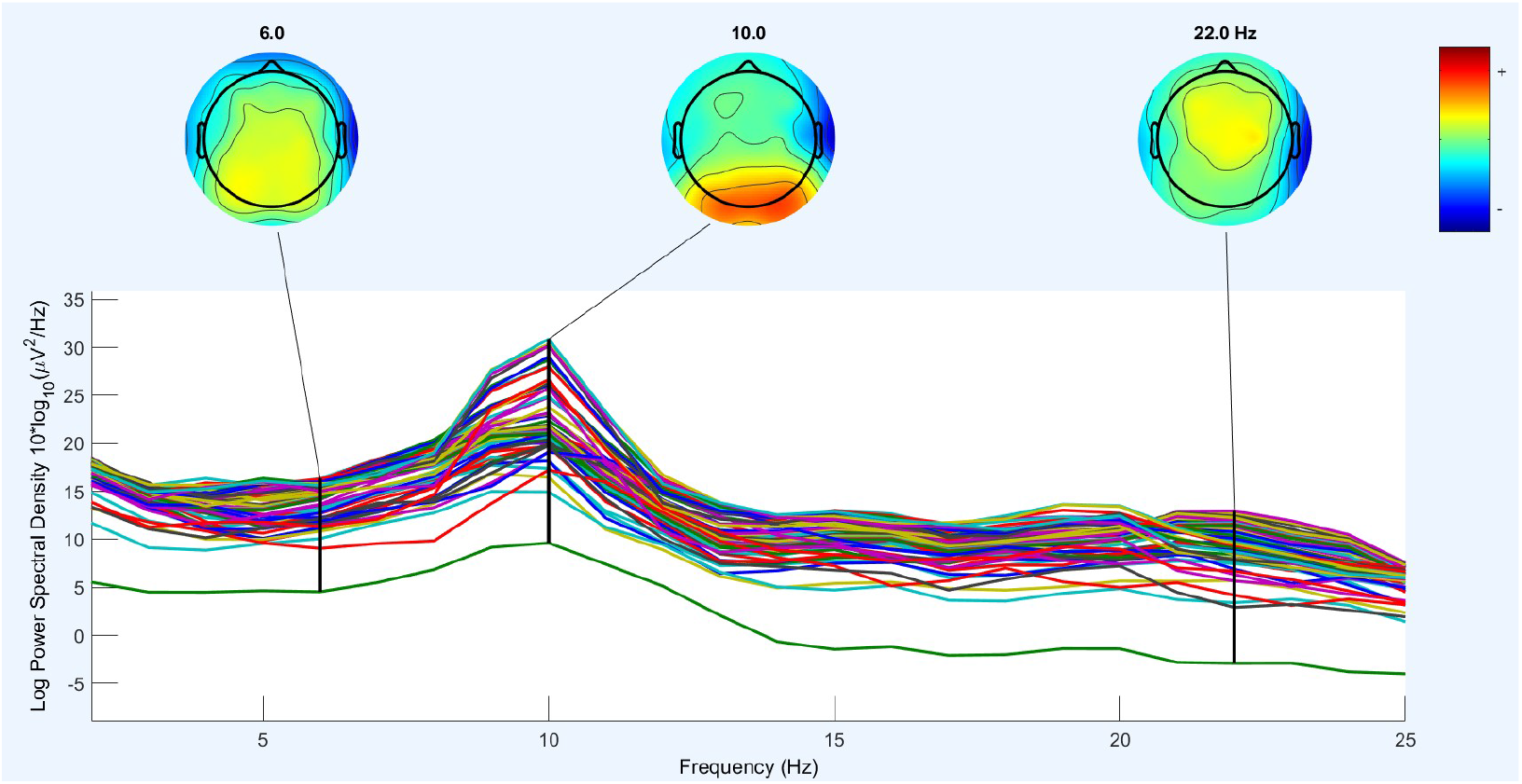
An illustration of channel spectrum. The lowest outlier is T10.

### II. Exploration of LP and LM interactions at coarser temporal scales: EEG trace segmented by 1.2 sec

It seems that the network interaction between LP and LM was more prominent at the finer temporal resolution. When the duration of the segments was extended from 0.4 sec to 0.8 sec and 1.2 sec, the correlation coefficients became less robust. The results of EEG segments of 0.4 sec and 0.8 sec are present in the main text.

**Table 1.**
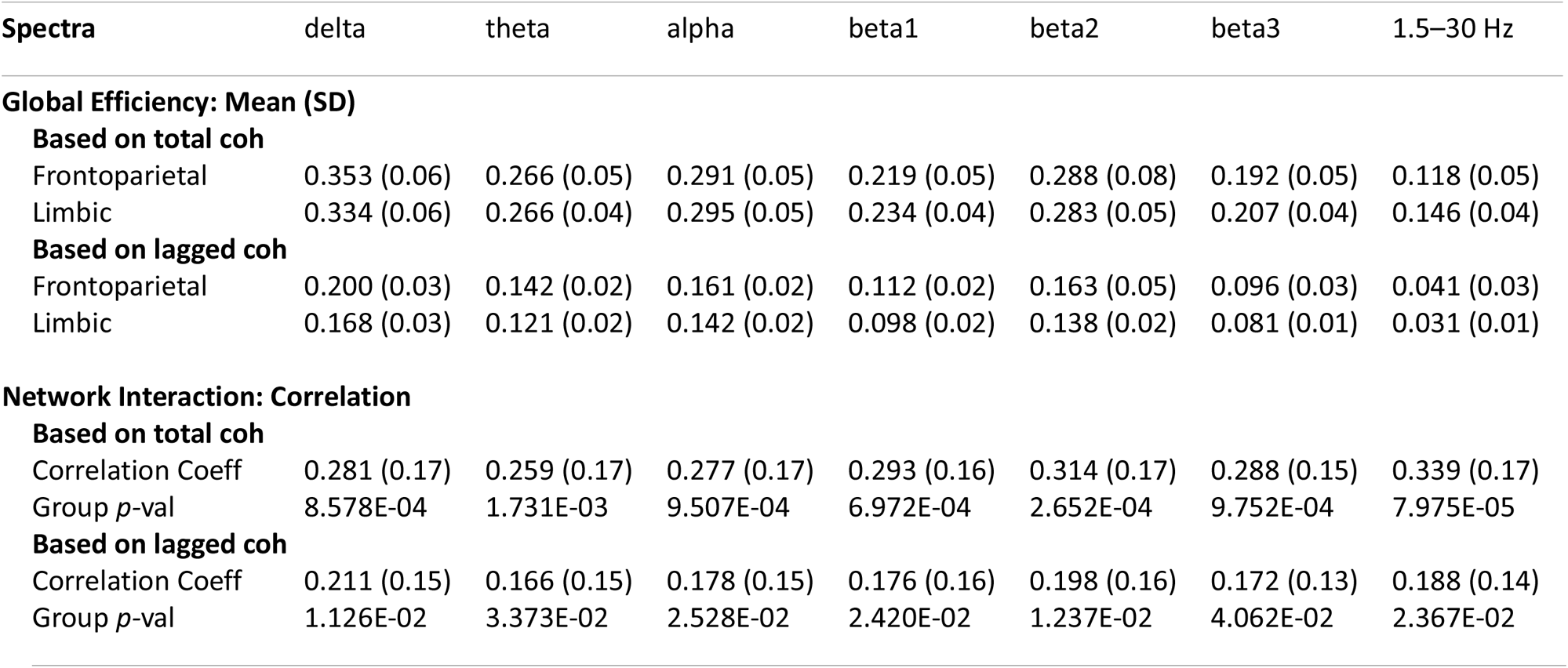
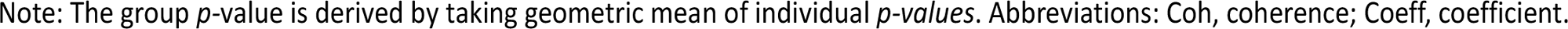
Global efficiency (upper) and its correlation (lower) between the frontoparietal and limbic networks (based on total and lagged coherence). EEG trace is segmented at the scale of 1.2 sec.

| Spectra | delta | theta | alpha | beta1 | beta2 | beta3 | 1.5–30 Hz |
| --- | --- | --- | --- | --- | --- | --- | --- |
| **Global Efficiency: Mean (SD)** |  |  |  |  |  |  |  |
| **Based on total coh** |  |  |  |  |  |  |  |
| Frontoparietal | 0.353 (0.06) | 0.266 (0.05) | 0.291 (0.05) | 0.219 (0.05) | 0.288 (0.08) | 0.192 (0.05) | 0.118 (0.05) |
| Limbic | 0.334 (0.06) | 0.266 (0.04) | 0.295 (0.05) | 0.234 (0.04) | 0.283 (0.05) | 0.207 (0.04) | 0.146 (0.04) |
| **Based on lagged coh** |  |  |  |  |  |  |  |
| Frontoparietal | 0.200 (0.03) | 0.142 (0.02) | 0.161 (0.02) | 0.112 (0.02) | 0.163 (0.05) | 0.096 (0.03) | 0.041 (0.03) |
| Limbic | 0.168 (0.03) | 0.121 (0.02) | 0.142 (0.02) | 0.098 (0.02) | 0.138 (0.02) | 0.081 (0.01) | 0.031 (0.01) |
| **Network Interaction: Correlation** |  |  |  |  |  |  |  |
| **Based on total coh** |  |  |  |  |  |  |  |
| Correlation Coeff | 0.281 (0.17) | 0.259 (0.17) | 0.277 (0.17) | 0.293 (0.16) | 0.314 (0.17) | 0.288 (0.15) | 0.339 (0.17) |
| Group *p*-val | 8.578E-04 | 1.731E-03 | 9.507E-04 | 6.972E-04 | 2.652E-04 | 9.752E-04 | 7.975E-05 |
| **Based on lagged coh** |  |  |  |  |  |  |  |
| Correlation Coeff | 0.211 (0.15) | 0.166 (0.15) | 0.178 (0.15) | 0.176 (0.16) | 0.198 (0.16) | 0.172 (0.13) | 0.188 (0.14) |
| Group *p*-val | 1.126E-02 | 3.373E-02 | 2.528E-02 | 2.420E-02 | 1.237E-02 | 4.062E-02 | 2.367E-02 |

Note: The group $p$ -value is derived by taking geometric mean of individual $p$ -values. Abbreviations: Coh, coherence; Coeff, coefficient.

**Table 2.** Average connectivity (upper) and its correlation between (lower) the frontoparietal and limbic networks across 7 spectra (based on total and lagged coherence). EEG trace is segmented at the scale of 1.2 sec.

| Spectra | delta | theta | alpha | beta1 | beta2 | beta3 | 1.5–30 Hz |
| --- | --- | --- | --- | --- | --- | --- | --- |
| **Average Connectivity: Mean (SD)** |  |  |  |  |  |  |  |
| **Based on total coh** |  |  |  |  |  |  |  |
| Frontoparietal | 0.386 (0.04) | 0.293 (0.04) | 0.318 (0.04) | 0.248 (0.05) | 0.333 (0.05) | 0.222 (0.05) | 0.161 (0.07) |
| Limbic | 0.402 (0.03) | 0.331 (0.04) | 0.344 (0.04) | 0.285 (0.05) | 0.359 (0.05) | 0.263 (0.05) | 0.212 (0.06) |
| **Based on lagged coh** |  |  |  |  |  |  |  |
| Frontoparietal | 0.154 (0.02) | 0.109 (0.01) | 0.123 (0.01) | 0.085 (0.01) | 0.130 (0.01) | 0.073 (0.01) | 0.027 (0.003) |
| Limbic | 0.140 (0.02) | 0.102 (0.01) | 0.114 (0.01) | 0.081 (0.01) | 0.117 (0.01) | 0.067 (0.01) | 0.025 (0.005) |
| **Network Interaction: Correlation** |  |  |  |  |  |  |  |
| **Based on lagged coh** |  |  |  |  |  |  |  |
| Correlation Coeff | 0.183 (0.14) | 0.173 (0.16) | 0.178 (0.17) | 0.172 (0.19) | 0.221 (0.21) | 0.182 (0.22) | 0.274 (0.26) |
| Group *p*-val | 3.968E-02 | 3.723E-02 | 2.422E-02 | 1.914E-02 | 3.725E-03 | 6.559E-03 | 2.197E-04 |
| **Based on total coh** |  |  |  |  |  |  |  |
| Correlation Coeff | 0.136 (0.14) | 0.116 (0.13) | 0.141 (0.14) | 0.157 (0.16) | 0.166 (0.15) | 0.155 (0.16) | 0.165 (0.17) |
| Group *p*-val | 9.275E-02 | 1.472E-01 | 8.931E-02 | 5.553E-02 | 4.852E-02 | 4.286E-02 | 3.655E-02 |

Note: The group *p*-value is derived by taking geometric mean of individual *p*-values. Abbreviations: Coh, coherence; Coeff, coefficient.

